# An extremes of phenotype approach confirms significant genetic heterogeneity in patients with ulcerative colitis

**DOI:** 10.1101/2021.10.04.462982

**Authors:** Sally Mortlock, Anton Lord, Grant Montgomery, Martha Zakrzewski, Lisa A. Simms, Krupa Krishnaprasad, Katherine Hanigan, James D. Doecke, Alissa Walsh, Ian C Lawrance, Peter A. Bampton, Jane M. Andrews, Gillian Mahy, Susan J. Connor, Miles P Sparrow, Sally Bell, Timothy H Florin, Jakob Begun, Richard B Gearry, Graham L. Radford-Smith

## Abstract

Ulcerative colitis (UC) is a major form of inflammatory bowel disease with increasing global incidence. There is significant phenotypic heterogeneity defined by a range of clinical variables including age of onset and disease extent. Clinical outcomes range from long-term remission on minimal therapy to surgical resection. Close to 70% of UC risk can be attributed to genetics and understanding the genetic mechanisms contributing to this risk and disease heterogeneity is vital for understanding disease pathogenesis and improving patient outcomes through targeted screening and therapies. This study aims to characterise the genetic heterogeneity of UC by identifying genomic risk variants specific to mild and/or severe forms of UC, exploring variations in the effect size of known risk variants and assessing the clinical value of a genetic risk score (GRS). We conducted genome-wide association (GWA) analyses in 287 patients with mild UC, 311 patients with severe UC and 583 age- and gender-matched controls. Odds ratios (OR) for mild vs control, severe vs control and combined mild and severe UC vs control were calculated. Using the combined UC data, two independent loci in the HLA region reached genome-wide significance. An additional genome-wide significant signal on chromosome 1 was identified in severe cases only. OR for known risk loci varied between mild and severe patients and were similar to previously published results. Effect estimates from the most recent UC GWA meta-analysis were used to calculate a GRS for each individual. A higher mean GRS was observed in both mild and severe UC cases compared to controls however, there was no difference between the mean GRS for mild and severe UC. Heterogeneity in effect sizes of UC associated variants between mild and severe disease burden suggests the presence of genetically distinct signatures. While large consortium data are needed to identify genome-wide significant variants, additional risk loci may be identified by targeted recruitment of individuals with a history of severe disease.

**Author Summary:** Ulcerative colitis (UC) is a chronic and often debilitating form of inflammatory bowel disease affecting approximately 0.3% of the population in industrialized economies. The disease displays significant clinical heterogeneity including age at presentation, disease severity, and the propensity to develop disease-related complications. Several previous studies have demonstrated the heritability of UC, identifying over 30 loci specific to the disease. The majority of these loci have small to modest effect sizes other than those within the Human Leucocyte Antigen (HLA) region on chromosome 6. Using stringent clinical criteria for defining mild and severe forms of UC in an extremes of phenotype approach, we undertook a genome wide association study in a dataset of 1222 participants to investigate genetic heterogeneity in this disease. We demonstrated substantial differences in genetic associations in severe UC as compared to mild UC. While over 2,000 SNPs achieved genome-wide significance in the severe UC analysis, none reached significance for mild UC. These results were reflected in significant differences in odds ratios. We identified Complement Factor B *(CFB)* as a potential susceptibility gene for severe UC in the Caucasian population with additional tissue gene expression demonstrating a positive correlation with disease severity.

## Introduction

Ulcerative colitis (UC) is a chronic inflammatory disorder of the large intestine and one of the major forms of Inflammatory Bowel Disease (IBD). IBD now has a global distribution and affects approximately 6.8 million of the world’s population^1^. Intestinal inflammation in UC is typically limited to the colonic mucosa and superficial submucosa. A number of factors have been implicated in contributing to disease severity including age at onset, disease extent, and genetic risk factors^2–6^. Individuals with mild disease may achieve adequate disease control through lifestyle modification and limited medical therapy such as the use of 5-aminosalicylates^7,8^. Those with moderate or severe disease are characterized by either severe attacks requiring hospitalization (acute severe UC; ASUC) and/or frequent disease flares that require corticosteroids, immunomodulators, and biologic drugs (chronic refractory UC). If intensive medical therapy fails to achieve sustained remission of symptoms, then surgery is regarded as a safe and effective option in achieving a reasonable quality of life^9,10^. The lifetime risk of ASUC is up to 25% and carries an additional increased risk of colectomy of up to 40% as compared to less than 15% in those individuals without a history of severe disease^11^. If we were able to predict disease severity at an early stage, more rapid escalation to advanced therapies may be instituted to attempt to change the natural history of the disease. Early identification of patients requiring more aggressive treatment options could assist in selecting the optimal treatment strategy on a patient-by-patient basis.

Prognostic factors that will assist both the patient and the treating team in predicting the course of the disease are the subject of several studies. Clinical risk factors that may assist in predicting risk of colectomy specifically at the time of diagnosis include extent of disease, age, need for systemic corticosteroids and either C-reactive protein (CRP) or Erythrocyte sedimentation rate (ESR)^3^. Factors that can predict future risk of ASUC at diagnosis include disease extent, CRP, and haemoglobin^12^. UC has an estimated heritability of 67% and the amount of variation captured by single nucleotide polymorphisms (SNPs) has been estimated as 33%^13,14^. The genetic basis of UC disease severity is informed by a limited number of studies that either focus on individual genes or regions such as the MHC^15,16^ and more recently GWAS and Immunochip studies^6,^. These and others^17-24^ have identified more than 120 independent loci associated with UC. Haritunians and colleagues developed a genetic predictor for UC refractory to medical therapy based upon a selection of 46 SNPs which included markers within the MHC^6^. Ten SNPs, all within the MHC region, reached genome-wide significance in their medical refractory case versus control analysis. An international study of IBD sub-phenotypes used a survival analysis to investigate markers associated with colectomy in UC. Five SNPs, all at 6p21 within the MHC, achieved genome-wide significance with the top SNP being rs4151651 (HR 1.72, 95% CI [1.47 – 2.00])^17^. This SNP is located in exon 5 of the Complement Factor B (*CFB*) gene on chromosome 6. *CFB* was also one of seven novel UC susceptibility genes identified in the first GWAS undertaken in the genetically distinct North Indian population^25^..

Monogenic mutations have been identified in specific IBD extremes of phenotype such as very early onset disease. However, these do not explain the majority of phenotypic variance in UC. Both mild and severe UC may represent polygenic conditions, sharing variants in the same genes that determine UC in the general population, or in genes novel to these extremes. In support of this, Lee and colleagues identified a single SNP intergenic between HLA-DRA and –DRB, rs9268877, that was associated with a poor UC prognosis (defined as need for anti-TNF therapy and/or colectomy) in a single centre study from Seoul, Korea^18^. Potential increases in statistical power afforded through analysis of extreme phenotypes has become an established approach to investigate complex disease^19,20^. Given the limited treatment options currently available for severe UC, in particular ASUC, there is an ongoing need to further define the genetic contribution to disease heterogeneity, to better understand severe disease pathogenesis, and identify novel and effective treatment targets. In this study we used a novel UC extremes of phenotype approach, carefully selecting criteria to define individuals with either severe UC or persistent, mild UC. The aims of the study are to further define the genetic differences between these subphenotypes and determine the value of a genetic risk score in predicting disease severity and hence UC outcome.

## Methods

### Patient samples and DNA isolation

Patients, and healthy controls, for this study were recruited from sites within the Australia and New Zealand IBD Consortium (ANZIBDC). Briefly, consecutive patients with a diagnosis of UC based on validated criteria^28^, were invited to join the ANZIBDC research program at each participating site. Phenotype data were based upon the Montreal classification^29^ together with additional detailed clinical data including smoking behaviour, medications, and surgery. Predetermined criteria were used to classify patients as either mild, or severe, UC.

Mild UC was defined as those individuals having a minimum disease duration and follow up of 10 years during which the patient was well-maintained on oral and/or rectal 5-amino salicylate therapy with oral corticosteroids limited to one course per 12 months, and with no history of corticosteroid dependence or intravenous corticosteroids. Patients with any history of immunomodulator therapy use of greater than 6 months and/or any biologic therapy were not considered as having mild UC. Severe UC was defined as those requiring colectomy due to: 1. Chronic active disease despite treatment with corticosteroids, an immunomodulator, and/or a biologic medication; and/or, 2. Acute severe disease having failed to respond to intravenous corticosteroids and/or rescue therapy with either infliximab or ciclosporin. Acute severe disease was defined by the Truelove and Witts criteria for all cases^30^. An additional 41 cases of acute severe UC, all satisfying the Truelove and Witts criteria, and who responded to rescue therapy with either infliximab or ciclosporin with persisting response to 12 months, were included in the combined UC cohort for all case-control analyses together with the mild and severe (colectomy) subgroups defined above and in Table 1.

**Table 1:** Cohort demographics. Numbers represent mean ±SD or absolute count (percentage) where appropriate. Percentages are calculated excluding missing data. Significance was calculated using either a Chi squared test or two sample t-test as appropriate.

|  | Control | Mild UC | Severe UC | P value |
| --- | --- | --- | --- | --- |
| **Demographics** |  |  |  |  |
| n | 583 | 287 | 311 |  |
| Female (%) | 337 (57.8) | 156 (54.4) | 146 (47.2) | 0.011 |
| **Smoking** |  |  |  |  |
| Ever (At Diagnosis) | 256 (44.4) | 73 (26.9) | 84 (29.7) | $1.032 \times 10^{-13}$ |
| At follow-up | - | 9 (6.7) | 7 (6.7) | 1 |
| Family History IBD (%) | - | 46 (25.7) | 34 (31.8) | 0.356 |
| **Disease Features** |  |  |  |  |
| Age at Dx (M±-SD) | - | 35.6 (15.1) | 32.6(14.2) | 0.001 |
| Disease duration (Years±-SD) | - | 20.4 (11.4) | 11.9 (10.1) | $< 2.2 \times 10^{-16}$ |
| Maximum disease extent (%) | | | | $< 2.2 \times 10^{-16}$ |
| E1 | - | 69 (24.0) | 3 (1.0) |  |
| E2 | - | 121 (42.2) | 70 (22.5) |  |
| E3 | - | 95 (33.1) | 233 (74.9) |  |
| Data not available | - | 2 (0.7) | 5 (1.6) |  |
| Colectomy | 0 (0.0) | 0 (0.0) | 311 (100.0) |  |
| Colectomy date (Years post Dx) | - | - | 6.4 (7.6) |  |
| Colectomy reason |  |  |  |  |
| Refractory disease | - | - | 157 (50.5) |  |
| Acute severe colitis | - | - | 148 (47.6) |  |
| CRC/dysplasia with refractory disease | - | - | 2 (0.6) |  |
| Data not available* | - | - | 4 (1.3) |  |
| **Treatment** |  |  |  |  |
| Anti TNF (%) | - | 0 (0.0) | 106 (37.4) |  |
| Adalimumab (%) | - | 0 (0.0) | 13 (5.7) |  |
| Infliximab (%) | - | 0 (0.0) | 83(37.2) |  |
| Other anti-TNF agent (%) |  | 0 (0.0) | 10 (3.2) |  |
| Cyclosporin (%) | - | 0 (0.0) | 63 (23.4) |  |
| 5ASA | - | 140 (96.6) | 71(76.3) | $4.5 \times 10^{-6}$ |
| Oral steroids (%) | - | 91 (50.6) | 113 (94.1) | $5.87 \times 10^{-15}$ |
| Immunomodulator (ever) | | | | $<2.2 \times 10^{-16}$ |
| Yes | - | 13 (4.6) | 241 (79.5) |  |
| *disease severity confirmed on histology |  |  |  |  |

A subgroup of the ulcerative colitis cohort from the lead site for this study (QIMR Berghofer MRI) underwent gene expression analysis for *CFB* using colonic tissue biopsies. Biopsies were collected by the principal investigator at the time of endoscopic examination. A total of 46 UC patients and 22 healthy controls were included in this analysis. Biopsies were taken from the sigmoid colon using a standard biopsy forceps technique, immediately snap frozen and stored at −80°C for RNA extraction, as previously described. Adjacent biopsies were taken from this segment for histological analysis. An inflammation score was generated for each biopsy site and each case using a validated scoring system^31^ (non-inflamed, n= 14; mild, n = 12; moderate, n = 16; severe n =4). RNA isolation and microarray analysis were performed as described below^30^.

Written informed consent was obtained from each patient as approved by the ethics committee of each member site. A blood sample was obtained from each participant. DNA isolation and quantification were performed using well-established protocols and as previously described.

### Genotyping

All genotyping was performed using Infinium technology (Illumina, San Diego, CA), specifically the OmniExpress chip containing 733,202 SNPs. Quality control (QC) was performed on genotypes using PLINK^33,34^. Call rates <0.95, SNPs with a mean GenomeStudio GenCall score <0.7, Hardy-Weinberg equilibrium P <10^−6^, and MAF <0.05 were excluded. Cryptic relatedness between individuals was identified by calculating a genomic relationship matrix in GCTA^35^. Ancestry outliers were identified using data from 1000 Genomes populations and principal components generated in GCTA. A total of 575,330 SNPs in 1,222 individuals remained for imputation. Genotypes were phased using ShapeIt V2 and imputed using the 1000 Genomes Phase 3 V5 reference panel on the Michigan Imputation Server^36^. Post-imputation QC was performed in PLINK removing imputed SNPs with low MAF <0.05 and poor imputation quality (R^2^ <0.8) leaving 6,273,901 autosomal SNPs for analysis.

### Data processing

#### Statistical analysis

GWAS analysis was performed for the combined UC cohort (639 cases and 583 controls), and mild (287 cases) and severe UC (311 cases) separately, using Logistic regression in PLINK. The first 5 principal components were used as covariates to account for population stratification and the genomic inflation factor was calculated (*λ*=1.02). Significant SNPs which survived the genome-wide correction (p<5×10^−8^) were cross checked against known SNPs for UC, Crohn’s Disease (CD) and combined IBD. A post-hoc analysis was performed restricting the number of SNPs tested to only 123 SNPs known to associate with UC from prior literature. Results for this analysis were considered statistically significant if a p-value <0.05 was obtained after Bonferroni correction including all tests from the combined group, mild only and severe only (k=369, critical α=1.36×10^−4^). Odds ratios for previously reported SNPs, which were significant in our dataset, were compared to investigate the consistency in effect sizes between studies and disease severity. Differences in odds ratios between mild and severe UC and between this cohort and published odds ratios were assessed using the Welch Modified Two-Sample t-Test.

To assess if the predictive power of SNPs differs between disease subtypes a genetic risk score (GRS) was calculated using summary statistics obtained from Liu et al^21^. Summary data based best linear unbiased prediction (sBLUP) was used to assign an effect size to each allele in the dataset based on the aforementioned summary statistics^35^. Individual GRSs were then calculated using the SNP effect estimates in PLINK. Two-sample t-testing was performed to test the association between GRS and disease by testing mild UC, severe UC and the combined UC cohort (n=639) against the control cohort independently. A further t-test between GRS of mild and severe UC was also performed.

GRS were also binned into deciles and the odds ratios of UC vs. control were calculated using the lowest decile as a reference. Sub-analysis using mild UC and severe UC vs. control were similarly performed. Further to the GRS, we calculated the risk score for medically refractory UC (scaled 0-92)^6^, which includes 46 equally weighted SNPs selected by Haritunians *et al*, however two of the SNPs were unable to be imputed. As such our score was modified to include the 44 remaining SNPs.

To investigate the association between the UC GRS and clinical factors, risk scores were regressed against disease extent (Proctitis, n=73; left-sided, n=207; extensive, n=352; total=634) and age at diagnosis (n=632) using the entire UC cohort. Differences in the mean disease extent and age of diagnosis between cases within the top and bottom 10% of risk scores was also tested. Associations between GRS and age were assessed as both a continuous variable and categorical (<20; 21-39; >40).

### Microarray analysis

Probes representing the *CFB* genes were obtained from dbSNP at NCBI (https://www.ncbi.nlm.nih.gov/snp). One previously reported SNP significantly associated with UC and for which we observe a much larger effect in severe UC, rs4151651, is a missense variant in an exonic region of complement factor B (*CFB*). To investigate the relationship between expression of *CFB* and UC severity we tested the association between *CFB* expression in the sigmoid and clinically diagnosed UC severity subgroups. Microarray gene expression data were read into R (version 3.4.1) using the Affy package version 1.56.0^37^. Probes were pre-processed using the expresso function where data were background corrected using the rma method quantile normalized and summarized using the median polish method. Data were filtered according to probe variance (cut-off: 0.5) and presence in all samples. Generalized linear regression was applied to identify a relationship between *CFB* expression and UC severity. P-values were adjusted using False Discover Rate (FDR). The probe 202357s was used as a proxy for CFB expression. One-way Analysis of variance with a Tukey’s post-hoc comparison between groups was applied to identify differences in the *CFB* probe between healthy controls and UC severity subgroups.

## Results

### Population

A total of 1222 participants were recruited for this study including patients with mild (n=287) and severe (n=352: n=311, colectomy and n=41, no colectomy) UC, as well as a matched healthy cohort (n=583) (Table 1). Control participants had a significantly higher prevalence of smoking compared to both the mild and severe UC subgroups (44.4% vs. 26.9% and 27.1%, respectively, P<0.001). Patients with severe UC were diagnosed younger than patients with mild UC (32.8 years vs. 35.6 years, P<0.01) and had a shorter disease duration (11.5 years vs. 20.4 years, P<0.001). As expected, there were significant associations between disease extent and disease severity. Specifically, limited disease (E1 or E2) was reported in 190 (66.7%) of mild UC patients compared to 90 (26%) of those with severe UC, while extensive disease (E3) was present in 257 (74.1%) of those with severe disease and only 95 (33.3%) of the mild UC subgroup (P<0.001). In contrast to previous studies^6^, family history of IBD was reported equally across both UC subgroups.

### Identified SNPs

A GWAS using the combined UC dataset identified 1,460 SNPs on chromosome 6 in the HLA region (lead SNP=rs28479879, OR=1.97, P=1.63×10^−14^) that were significantly associated with UC reaching a conventional genome-wide significance threshold of P<5×10^−8^ (Supplementary Figure 1a). Conditioning on the lead SNP in this region identified a secondary independent risk locus in the HLA region (lead SNP=rs144717024, OR=5.52, P=1.57×10^−10^). When considering only patients with severe UC, 2,018 SNPs were significantly associated, including a locus on chromosome 1 (lead SNP=rs111838972, OR=1.82, P=6.28×10^−9^) near *MMEL1* and a locus in the HLA region on chromosome 6 (lead SNP=rs144717024, OR=12.23, P=1.7×10^−19^) (Supplementary Figure 1b). Conditioning on the lead SNP in each of these regions identified a secondary independent risk locus in the HLA region (lead SNP= rs6916742, OR=2.18, P=1.41×10^−10^). The risk loci also pass a more stringent Bonferroni correction (P<7.97×10^−9^) accounting for the total number of SNPs tested. The large effects observed for variants in the HLA region are consistent with previous reports of large effects of UC associated haplotypes in this region^38,39^. Effect sizes observed were substantially reduced when considering mild UC patients only, resulting in no significant SNPs reaching genome-wide significance when compared to control participants. However, the direction of these effects was consistent with severe UC. The OR for the lead SNPs, rs28479879 (lead SNP in combined), rs144717024 (lead SNP in combined and severe) and rs111838972 (lead SNP in severe), were significantly higher in severe UC compared to mild UC (P <9×10^−6^).

When comparing our results to the 123 previously identified SNPs associated with UC we were able to replicate seven SNPs in our dataset (Table 2). Of the 123 previously identified SNPs tested, 55% (n=68) had larger effects in severe UC cases compared to mild cases (Table 2). We do, however, observe large standard errors for OR estimates in this study due to the relatively small sample size. Overall, a large proportion of SNP effects were in the same direction as those reported previously (88% combined UC, 82% mild UC, 85% severe UC). One SNP, rs7554511, on chromosome 1 was only associated with mild cases and not severe, or combined, UC cases. rs4151651 had a statistically higher OR in severe UC compared to mild UC (P=1.08×10^−31^). Similarly, the ORs for three SNPs estimated in the combined UC cohort and severe UC cohort were significantly different from the published estimates (rs4151651 P_combined_=8.06×10^−16^, P_severe_=2.56×10^−55^; rs6667605 P_combined_=2.31×10^−4^, P_severe_=8.70×10^−10^; rs10761648 P_combined_=8.57×10^−4^, P_severe_ =1.99×10^−3^) (Table 2; Figure 1). In all three cases the published OR was most similar to the mild UC OR estimate.

**Table 2:** Seven published SNPs associated with ulcerative colitis (UC) replicated in association analyses for combined UC cases and mild and severe cases only.

| Previously Identified SNPs |  |  |  |  |  |  | Combined cases |  | Mild Cases |  | Severe Cases |  |
| --- | --- | --- | --- | --- | --- | --- | --- | --- | --- | --- | --- | --- |
| SNP | CHR | BP | Effect Allele | OR | P.value | Reference | OR<br>(95% CI) | P.value | OR<br>(95% CI) | P.value | OR<br>(95% CI) | P.value |
| rs6667605<sup>+</sup> | 1 | 2502780 | C | 1.09 | 3.16E-10 | 19 | 1.39<br>(1.19-1.64) | 5.13E-05** | 1.14<br>(0.94-1.39) | 1.97E-01 | 1.72<br>(1.41-2.13) | 1.00E-07** |
| rs80174646 | 1 | 67708155 | G | 1.61 | 4.34E-62 | 19 | 2.04<br>(1.43-2.94) | 1.01E-04** | 2.17<br>(1.33-3.57) | 1.74E-03 | 1.92<br>(1.23-3.03) | 4.57E-03 |
| rs7554511 | 1 | 200877562 | C | 1.18 | 6.83E-31 | 19 | 1.39<br>(1.16-1.67) | 4.28E-04 | 1.61<br>(1.273-2.04) | 9.56E-05** | 1.21<br>(0.96-1.51) | 1.02E-01 |
| rs4151651<sup>+\$</sup> | 6 | 31915614 | A | 1.72 | 6.05E-12 | 17 | 3.73<br>(2.48-5.63) | 2.72E-10 | 1.81<br>(1.07-3.06) | 2.79E-02 | 6.00<br>(3.86-9.35) | 2.03E-15 |
| rs9268853 | 6 | 32429643 | T | 1.41 | 1.35E-55 | 21 | 1.92<br>(1.59-2.33) | 4.11E-12** | 1.54<br>(1.23-1.92) | 1.57E-04* | 2.44<br>(1.89-3.13) | 9.10E-13** |
| rs10761648<sup>+</sup> | 10 | 64354262 | T | 1.16 | 2.99E-15 | 24 | 1.53<br>(1.24-1.89) | 7.86E-05** | 1.46<br>(1.13-1.89) | 3.82E-03 | 1.57<br>(1.22-2.02) | 5.03E-04 |
| rs2836878 | 21 | 40465534 | G | 1.25 | 7.35E-53 | 19 | 1.43<br>(1.18-1.7) | 2.92E-04* | 1.47<br>(1.15-1.89) | 1.81E-03 | 1.35<br>(1.06-1.72) | 1.18E-02 |
| *Significant association accounting for multiple testing in individual association analysis<br>**Significant association accounting for multiple testing across all three analyses<br><sup>+</sup> Published OR estimates are significantly different from combined and severe GWAS estimates.<br><sup>\$</sup> OR estimates are significantly different between mild and severe GWAS. | | | | | | | | | | | | |

**Figure 1.**
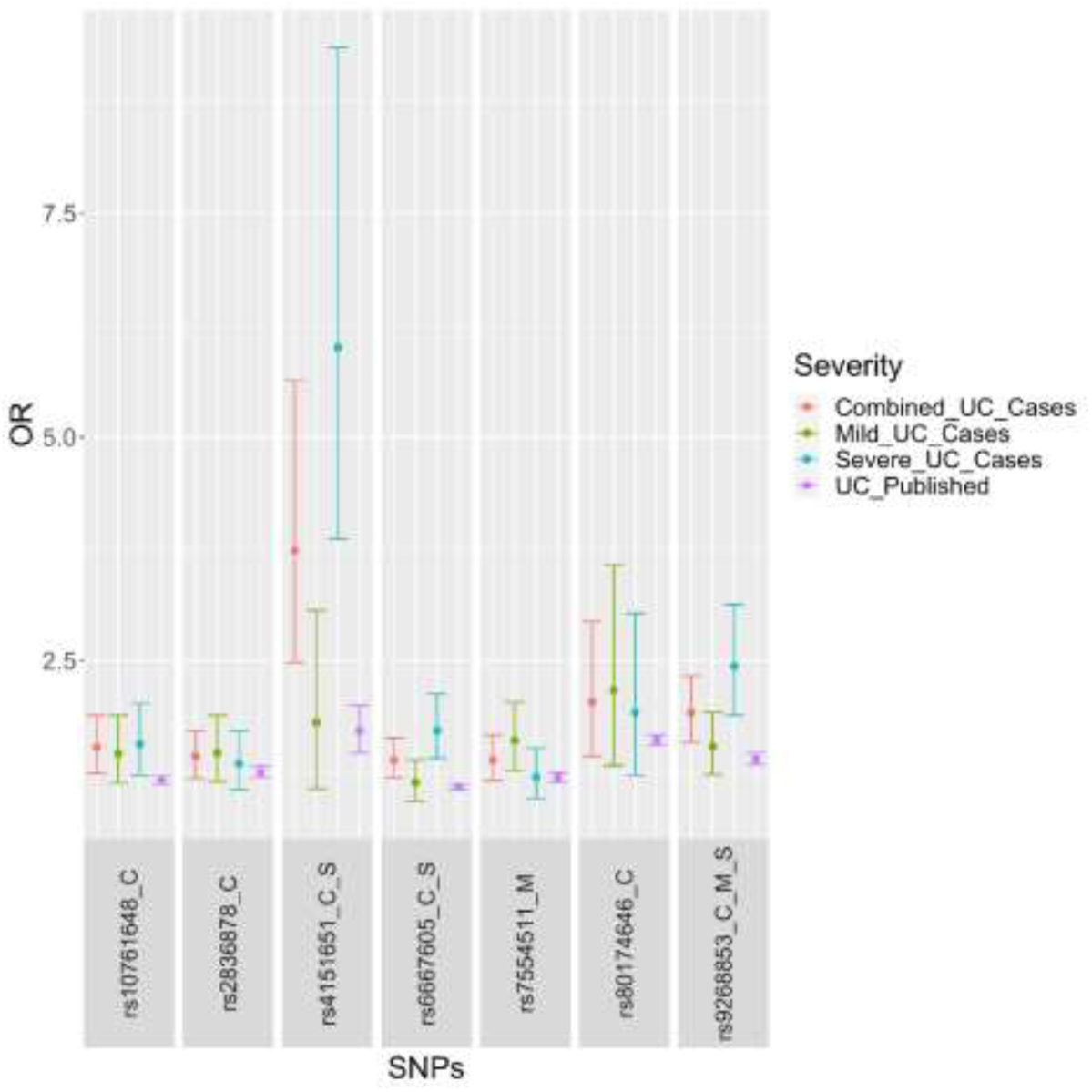
Odds ratios with 95% confidence intervals for seven published SNPs associated with Ulcerative Colitis (UC) and replicated in association analyses for combined UC cases (C) and mild (M) and severe cases (S) only.

### Genetic risk score

Genome wide risk scores were significantly increased in both mild (P=9.60×10^−13^) and severe UC compared to controls (P=8.03×10^−16^), however, no difference between mild and severe UC was observed (Figure 2). Considering all UC patients as a single group vs controls, the genome-wide risk score was also significantly higher (P<2.2×10^−16^). When separated into deciles (Figure 3), the proportion of control participants reduced from 79.8% (decile 1) to 28.7% (decile 10) as the genetic risk score increased. Conversely the proportion of severe patients increased from 9.7% (decile 1) to 34.4% (decile 10) with increasing risk score. Similarly, the proportion of patients with mild UC increased from 10.5% (decile 1) to 32.8% (decile 10). Odds ratio calculations between the lowest and highest deciles showed an increased proportion of participants in the highest decile had UC (either mild or severe) compared to the lowest decile (OR=9.18, 95%CI=5.12-16.47, Z=7.3, P=1×10^−4^). There was a significant positive association between UC GRS and disease extent (P=4.91×10^−3^) and a significant difference (P=0.023) in disease extent between cases in the top and bottom deciles. Age at diagnosis was not significantly associated with the GRS when assessed as either a continuous or categorical variable.

**Figure 2:**
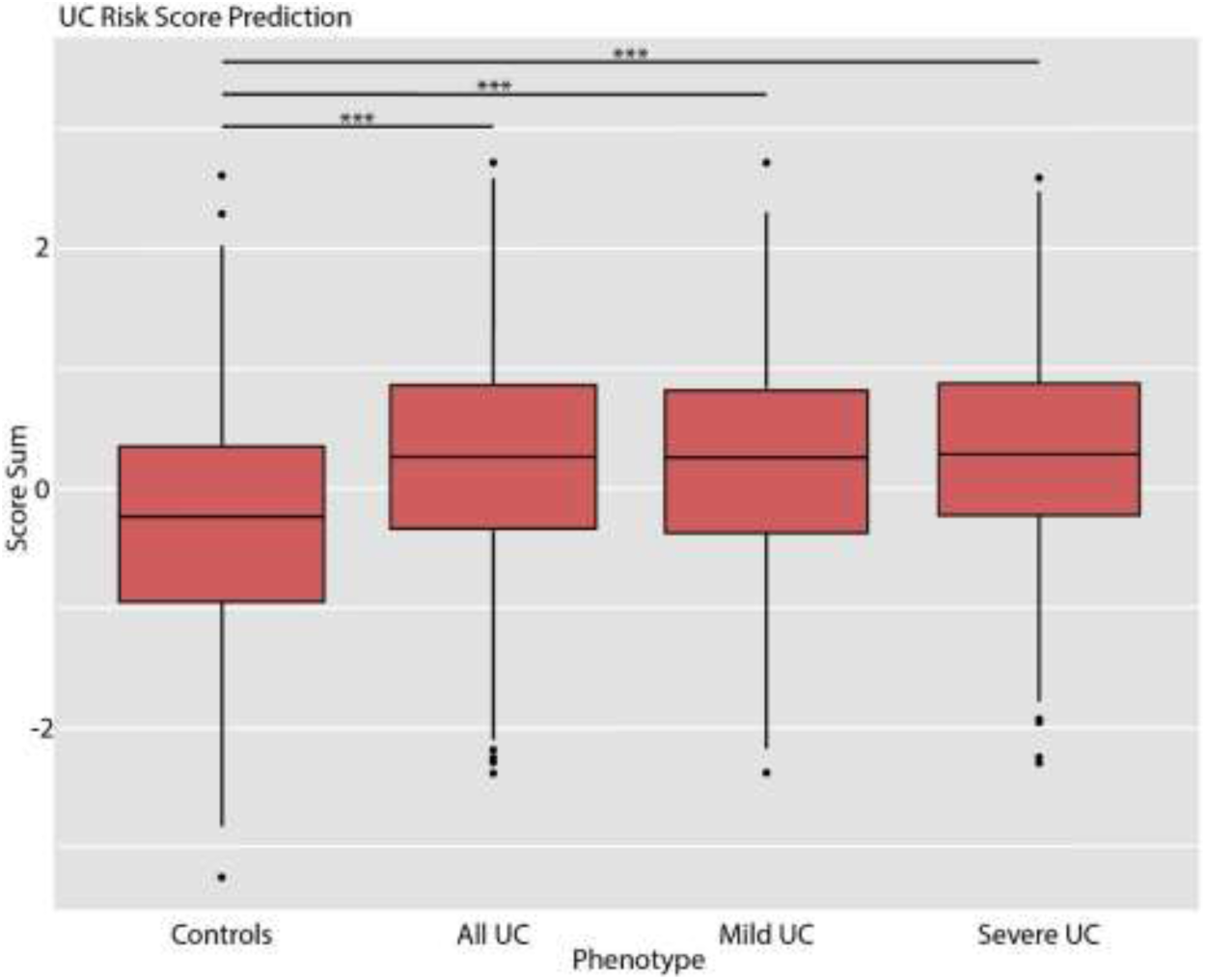
Distribution of UC genetic risk scores for control patients (Controls), both severe and mild UC patients (All UC), mild UC patients only (Mild UC) and severe UC patients only (Severe UC).

**Figure 3:**
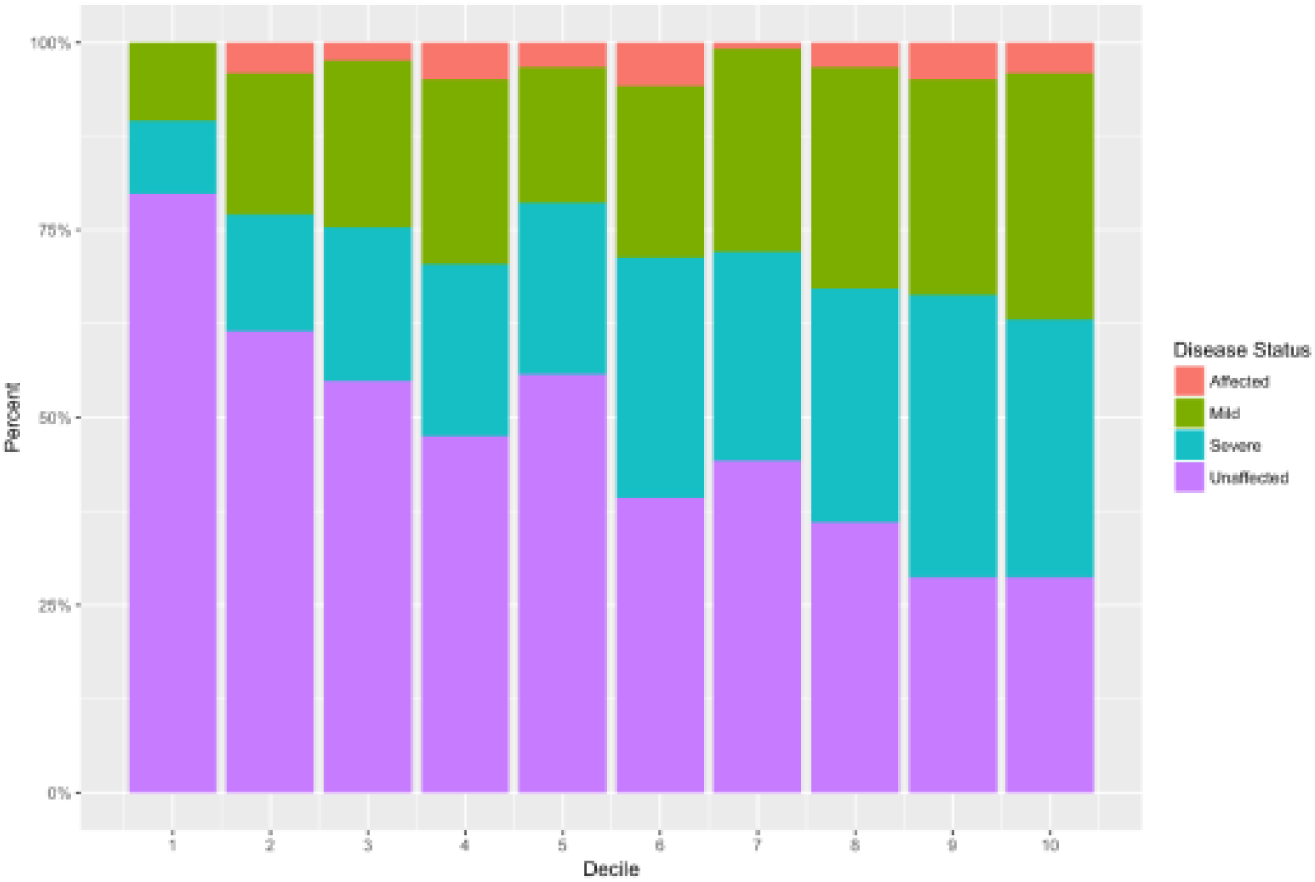
Patients divided into deciles according to UC genetic risk score and the proportion of patients with mild (green), severe (blue), severe without colectomy (red) and unaffected (purple).

No significant association was observed between the previously published medically refractory UC risk score^6^ and our population (P=0.318). No significant difference in the proportion of mild and severe UC in the highest and lowest deciles was observed (OR=1.25, 95%CI=0.52-3.01, Z=0.498, P=0.619). Furthermore, a post-hoc analysis did not reveal any significant increase in risk scores^6^ of either our medically refractory UC (P=0.57), or our acute severe UC (P=0.59) subgroups, when compared to control subjects (Figure 4).

**Figure 4:**
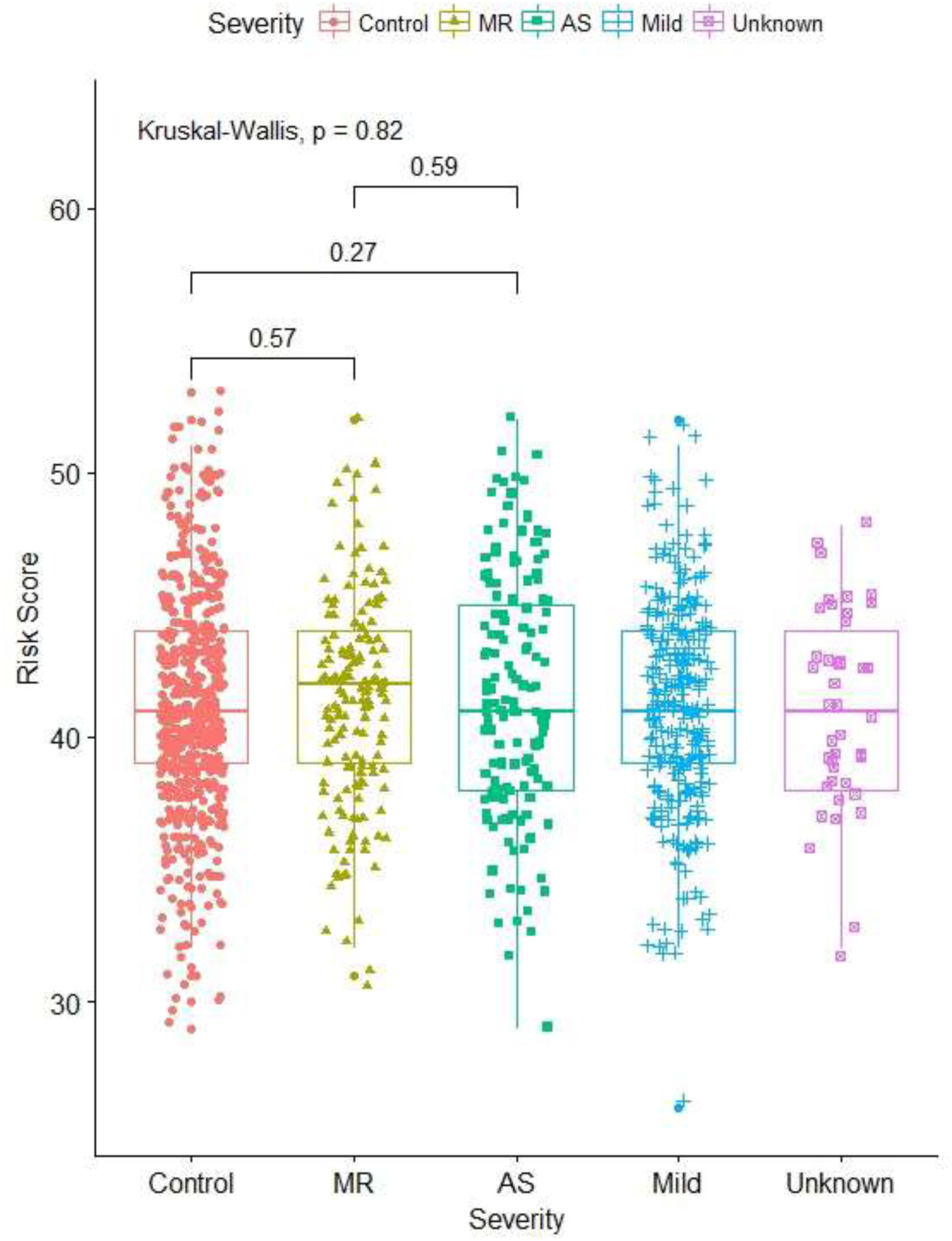
Medically refractory UC risk scores calculated using the formula from Haritunians et al., for control, medically refractory (MR), acute severe (AS), mild UC subjects and those with acute severe UC without colectomy (AS responder).

Using the AVENGEME R package^38^ we estimate that a training set of ∼22,000 individuals would be required to achieve a clinically relevant AUC of 0.75 using 100,000 SNPs if the genetic variance explained is 33% (SNP heritability) and the proportion of SNPs having no effect on disease is 0.90 (Supplementary Table 1).

#### *CFB* gene expression

Regression analysis indicated an increase in *CFB* expression in sigmoid colon mucosa in the UC patients (p = 0.002, FDR = 0.037). The expression of *CFB* was significantly different between the control group and mild UC and between the control group and moderate UC (Figure 5, Tukey’s test, p < 0.0001). In contrast, *CFB* expression in UC non-inflamed sigmoid was similar to healthy controls (Tukey’s test, p = 0.25).

**Figure 5:**
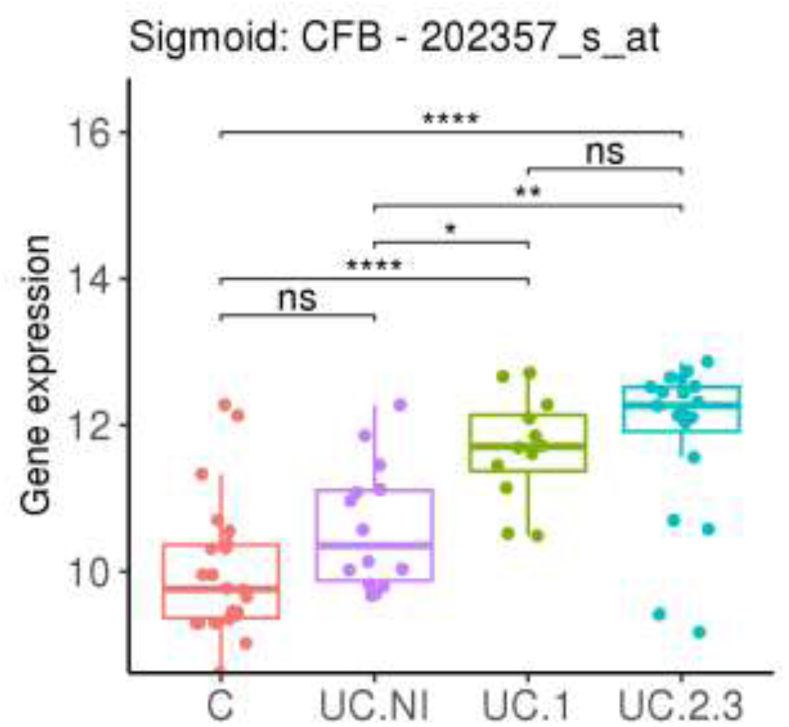
Microarray gene expression levels for *CFB* using probe 202357_s_at, for controls (C), non-inflamed UC (UC.NI), mild UC (UC.1), and moderate to severe UC (UC.2.3).

## Discussion

Genome-wide association studies, using large international cohorts, have identified over 200 SNPs linked to IBD that explain approximately 8.2% of the variance in UC risk^21,23,25^. These studies have been invaluable in identifying SNPs that explain disease susceptibility and hence provide important insights into disease pathogenesis. However, these SNPs do not differentiate between patients who experience particularly aggressive forms of UC as opposed to those with persistent, documented, mild UC. Without the granularity of data to separate these sub-phenotypes, genetic influences reported in the literature to date may provide only part of the unique genetic signatures carried by each form of UC. In this study we assess two distinctly different groups of patients with UC, namely those who follow a severe course which typically requires surgery within a median of 6.4 years from diagnosis and those who have been diagnosed and followed up for at least 10 years with limited medical interventions required to control disease activity and no requirement for surgery. Previous studies indicate that these two extremes of UC phenotype account for between 25 and 40% of all UC cases^2,3,9-11^.

Our study finds the effect sizes of known UC risk variants differ between patients with severe UC and mild UC. Notably, only one SNP was identified, rs7554511, which was related to mild but not severe UC in our dataset. Effect sizes reported in this study are on average 7% larger than in the published literature. This effect was even more pronounced when considering only patients with severe disease (10%). Even our mild UC subgroup had an effect size comparable with published effect sizes, suggesting international meta-analyses may use a mixture of patients with severity typically on the milder side of the disease spectrum. This may relate to the recruitment process for genetic studies with many patients identified from outpatient clinics and population-based registries. The observations for mild UC are supported by those of Kopylov and colleagues^41^. In a North American IBD Consortium analysis of 156 index SNPs from known IBD loci in their mild UC cohort, none achieved the pre-defined significance threshold.

For severe UC, of note is rs4151651, a SNP in an exonic region of complement factor B (*CFB*). This SNP had a much larger odds ratio (6.00) in patients with severe UC compared to mild UC (1.81). CFB is a secreted protein in the alternative complement pathway and is mainly expressed by mononuclear phagocytes. The complement system plays important roles in pathogen recognition and clearance^42^, and both inflammatory and immune responses. It has also been implicated in a range of autoinflammatory disorders including IBD^43^. Recent multi-ethnic studies in IBD genetics have identified *CFB* as one of two novel UC susceptibility genes in the North Indian population, with *CFB* allelic heterogeneity demonstrated when comparing North Indian, Japanese and Dutch populations^27,44^. The driver SNP, rs537160, in the UC associated Dutch haplotype was also replicated in this study in the combined (P=2.48×10^−5^) and severe (P=2.07×10^−9^) GWAS, and was a predicted transcription factor binding site for POLR2A and TFAP2A^44^. The over representation of the rs4151651 and rs537160 risk alleles in patients with severe UC may be associated with abnormal complement factor B secretion, impaired pathogen clearance within the colonic mucosa, and/or an exaggerated and poorly controlled immune response. Our gene expression data support a potential role for *CFB* in the mucosal inflammatory response typical of severe UC with a stepwise increase in expression across a spectrum of disease activity from remission through to severe disease. These observations replicate and extend previous *CFB* gene expression analysis in the context of UC^43^. The study by Ostviks and colleagues identified the colonic epithelium as the major local source of this increased *CFB* expression in active UC. Functional analysis of a SNP (rs12614) in *CFB* demonstrated significantly reduced alternate complement pathway activity in UC sera from individuals homozygous or heterozygous for this variant as compared to homozygous wild-type^27^. Whilst rs12614 is not in LD with rs4151651 or rs537160 it suggests a possible role for genetic regulation of *CFB* in UC. Studies in animal models of IBD have identified potential pathogenic and protective roles for different Complement pathway components in disease aetiology. Specifically, an alternative pathway knockout ameliorated the early effects of a dextran sodium sulphate-induced colitis^45^, and subsequent work demonstrated therapeutic potential for CR2-fH, a targeted inhibitor of the alternative pathway^46^. There has also been interest in the development of agents that can block complement pathway components such as C5a or its receptor. The far stronger association with severe UC in this study supports genetic heterogeneity within UC and the need to further explore the genetic regulation of Complement in mucosal immune responses and how this is influenced by local environmental factors such as the intestinal microbiome.

In our study, people in the highest decile of the genetic risk score are 9 times more likely to have UC compared to those in the lowest decile of genetic risk. We also found a significant association between disease extent and the genome-wide GRS developed on all UC calculated in this study. However, the GRS was unable to separate mild, from severe, UC in our cohort. This limitation to the GRS based upon currently available data likely reflects the milder disease course of many UC participants in GWAS studies to date and the clinical data available to define extreme phenotypes. There may be a lack of access to patients who have undergone surgery for severe UC given that their follow up is often with the surgical service at their local hospital, and that they remain a minority within the total recruited UC population. As such, independent larger and well-defined subgroups would be required to further develop robust indicators of disease course.

To date, two publications have explored genetic nuances between patients with mild and severe forms of UC^6,18^. In the first of these, Haritunians *et al*., found that medically refractory UC was associated with extensive disease, family history and 46 SNPs. When using 44 of the 46 SNPs identified by Haritunians^6^ to calculate a GRS, we found no association with disease severity. Our study used a stricter definition of mild UC, specifically, all patients in this subgroup had not undergone colectomy within 10 years of diagnosis, had not experienced an episode of severe colitis requiring hospital admission and/or intravenous corticosteroids nor required immunosuppression therapy for greater than 6 months. These extremes of phenotype criteria are similar to those used by Lee and colleagues in their analysis of a Korean UC cohort, and likely result in more distinct mild, and severe, UC subgroups^18^. This study of UC identified one SNP that was associated with the severe subgroup and which reached genome wide significance. This SNP, rs9268877, was not associated with overall UC disease susceptibility.

The strengths of our study include the *a priori* case definitions for mild, and severe, UC, the recruitment of population controls from the same population, and the detailed clinical metadata ascertained for all cases. Clinical and genetic findings are predominantly consistent with previous published data while highlighting the genetic heterogeneity within the sub-phenotype of UC. Limitations relate to statistical power across the study and within subgroups.

## Conclusion

Mild and severe forms of UC show distinct genetic signatures characterised by differences in effect sizes of risk variants. Genetic heterogeneity between sub-phenotypes can make the development of a diagnostic genetic risk score difficult. While the direction of effects is relatively consistent, the influence of genetics on mild UC is noticeably reduced with no statistically significant hits at the genome-wide significance level in our dataset. Combining mild and severe patients into a single cohort for GWAS increases genetic heterogeneity, likely reducing the ability of the GRS to distinguishing between clinically relevant sub-phenotypes. We identified *CFB* as an important candidate for UC susceptibility within a Caucasian population and highlighted its potential role in determining UC severity. Future studies should consider the severity of disease when trying to elucidate genetic nuances of UC.

## Acknowledgments

We wish to acknowledge funding for this work from the following: National Health and Medical Research Council project grant funding; QIMR Berghofer MRI laboratory support funding; RBWH Foundation grant funding. We are grateful to all the participants who took part in the study and to the clinical nurses, administrative staff, and research nurses who assisted in the study.

---

**Supplementary Table 1.** Estimated number of cases and controls (in 1000s) required to achieve a clinically relevant AUC using 1,000,000 SNPs that explain half the heritability of liability of Ulcerative Colitis given a disease prevalence of 0.0013, heritability 0.67 and 1:1 ratio of cases and controls.

| AUC | Proportion of null SNPs |  |  |  |
| --- | --- | --- | --- | --- |
|  | 0.99 | 0.90 | 0.75 | 0 |
| 0.75 | 4 | 22 | 34 | 35 |
| 0.80 | 5 | 32 | 56 | 63 |
| 0.85 | 8 | 52 | 98 | 129 |
| 0.90 | 18 | 127 | 254 | 411 |

**Supplementary Figure 1:**
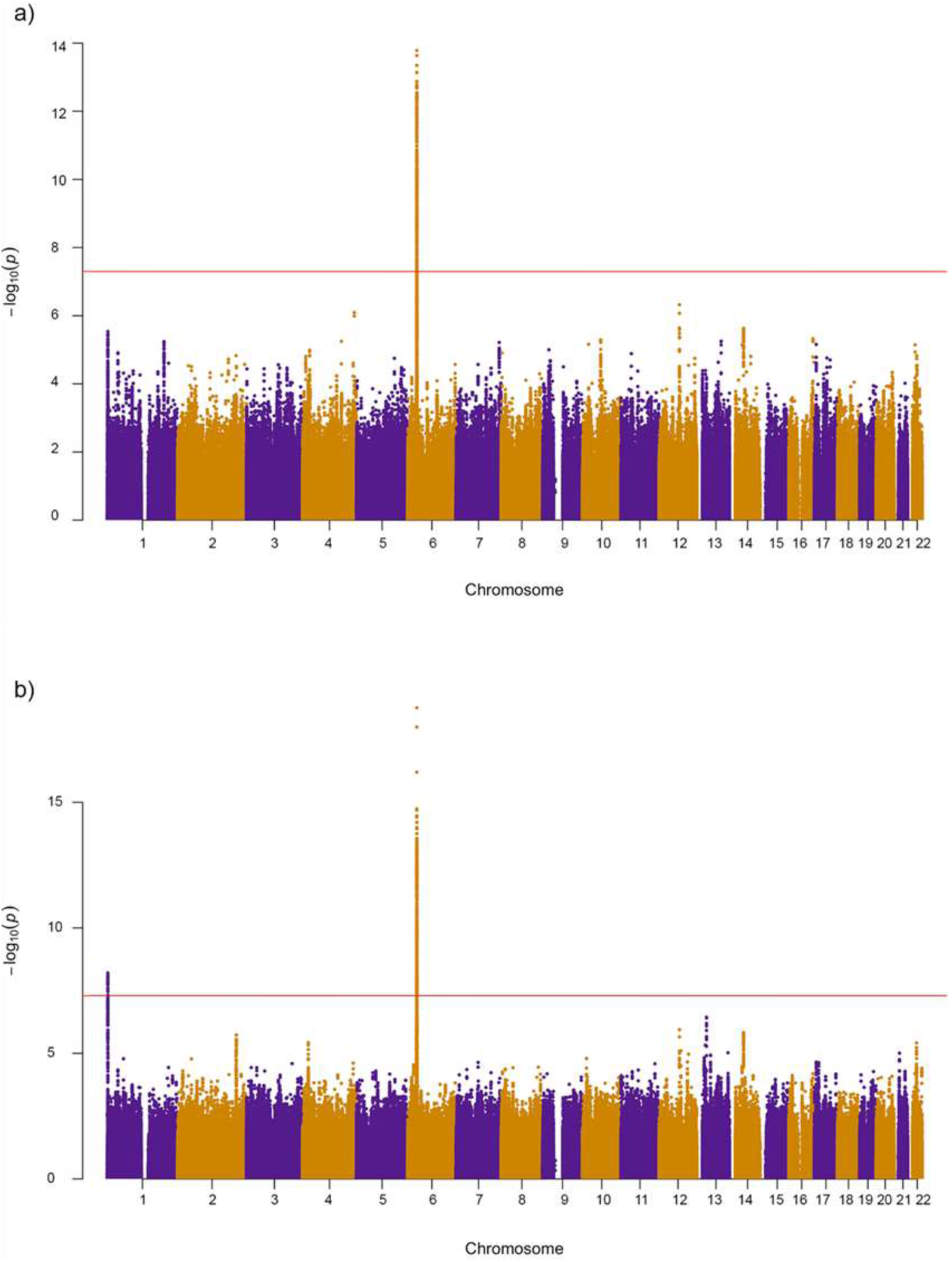
Manhattan plots for **(a)** mild and severe UC patients groups combined vs healthy controls and **(b)** severe UC patient group only vs healthy controls.

